# Activation of thalamo-cortical circuits with posterior hypothalamic nucleus deep brain stimulation

**DOI:** 10.1101/2021.05.16.444334

**Authors:** Calvin K. Young, Brian H. Bland

## Abstract

The posterior hypothalamic nucleus (PH) has extensive anatomical connections to motor, cognitive, visceral, and homeostatic areas of the brain and serves as a crucial subcortical modulator of behaviour. Previous studies have demonstrated that deep brain stimulation (DBS) of this area can lead to powerful activation of motor behaviour, overcoming two rodent models of parkinsonian akinesia by increasing neocortical excitability. However, it is unclear how the PH may mediate this increase in neocortical excitability. In the present study, we examined the role of the thalamus in the PH-DBS mediated increase in neocortical excitability. In urethane anaesthetized animals, we demonstrate that PH-DBS elicits increased spiking activity in the motor thalamus (VL) that receives direct afferents from the PH that precedes the increase in spiking activity in the corresponding motor cortex. In contrast, in the somatosensory thalamus (VPM) where PH afferents are sparse at best, PH-DBS did not elicit an increase in thalamic activity despite of a slight increase in the corresponding somatosensory cortical spiking. Current source density analyses suggest a thalamo-cortical mechanism for motor cortex activation whereas a cortico-cortical activation mechanism is involved in somatosensory cortical activation. Inactivation of the VL resulted in the abolition of motor cortex spiking despite of the persistence of desynchronized field potential activity. Collectively, these data suggest indirect orthodromic activation of PH output fibres to the thalamus mediates increased neocortical excitation, which may spread through cortico-cortical connections and lead to an increase in integrated, non-stereotypical motor behaviour.

---

The ability for chemical or electrical stimulation of the posterior hypothalamic nucleus (PH) to elicit locomotion has been well-documented (Bland and Vanderwolf, 1972; Shekhar and DiMicco, 1987; Lammers et al., 1988; Marciello and Sinnamon, 1990; Sinnamon et al., 1991; Slawinska and Kasicki, 1995; Oddie et al., 1996; Slawinska and Kasicki, 1998). Although the nature of PH-elicited locomotion is controversial (Shekhar and DiMicco, 1987; Lammers et al., 1988), it is clear that at the behavioural level it is not a mere direct activation of brainstem and spinal pathways that increases the magnitude of motor behaviours, but an integrated response where rats can engage in a whole repertoire of motor behaviours such as jumping, bi-directional turning, rearing and climbing (Bland and Vanderwolf, 1972).

While there is anatomical and electrophysiological evidence that the PH does directly modulate motor systems at the brainstem and spinal levels (Murphy and Gellhorn, 1945; Granit and Kaada, 1952; Kausz, 1990; Vertes and Crane, 1996), little attention has been paid to how a complex array of motor behaviours may be coordinated involving higher-order brain areas. Past research has shown the inactivation of the septal complex leads to hypokinesia, and blocks PH-elicited wheel-running behaviour (Mizumori et al., 1989; Lawson and Bland, 1993; Oddie et al., 1996). Since wheel running is a relatively specific behaviour that does not tap into the rich complexity of motor acts demonstrated by an animal in more freeform tasks such as the open field, and that the long-term consequence of septal lesion is hyperkinesia (Gray and McNaughton, 1983), it remains unclear what role the septum plays in PH-elicited locomotion.

The previous study has demonstrated that PH stimulation increases blood oxygen-dependent level responses mostly in the neocortex and the periaqueductal gray (Young et al., 2011b). Sparse activation was seen in the thalamus but was robust under high-intensity stimuli. Electrophysiological recordings from a sample of activated neocortical areas suggested a general increase in spiking, coupled with cortical desynchronization. These changes in cortical excitability were also shown to decrease movement thresholds for movements elicited from cortical stimulation. These data collectively suggest the complexity of motor behaviours exhibited by animals with PH stimulation may be at least partly due to the activation of cortical circuitry. It is, however, not clear how this increase in cortical excitability was mediated. There are direct reciprocal connections found between the PH and some of the cortical areas activated by it, such as the motor and cingulate cortices (Vertes et al., 1995; Cavdar et al., 2001). The previous study found no evidence for antidromic activation, and the sparse cortical afferents from the PH are unlikely to support extensive activation seen in imaging and electrophysiology studies. Given the extensive, unidirectional projections from the PH to the thalamus (Vertes et al., 1995), it is likely that cortical excitation is mediated through various thalamic nuclei.

Intriguingly, the PH does not project to all nuclei of the thalamus. For example, the PH provides modest innervations to the laterodorsal (LD) and ventrolateral (VL) nuclei of the thalamus, but none to the ventroposterior medial nucleus (VPM; Vertes et al., 1995). In the present study, the role of the thalamus is examined in the context of PH-mediated increase in cortical excitability. Paired thalamic and cortical recordings with different PH stimulation protocols were made to assess: 1) thalamic responses to PH stimulation; 2) temporal relationships between thalamic and cortical activities and; 3) the effect of changing PH stimulation parameters on thalamic activity. Additionally, thalamic inactivation by the infusion of procaine, a local anaesthetic, was performed to elucidate the dependence of cortical spiking on the thalamus. Given previous findings, it was hypothesized that PH stimulation increased cortical excitability indirectly through activating the thalamus. Data reported here supports the above hypothesis, and further suggests the modulation of thalamo-cortical excitability is pathway-dependent, affecting only those thalamo-cortical systems receiving PH-thalamic projections.

## Materials and Methods

### Recording Procedures

Fourteen male Long Evans rats were obtained from the Animal Care Facility at the University of Calgary. The animals were given food and water *ad libitum* under 12/12 light/dark cycles. Rats received halothane anaesthesia (1.5% oxygen alveolar concentration) for the implantation of a jugular vein cannula used to administer the anaesthetic ethyl carbamate (1.2 g/kg) for the rest of the experiment. The rats were monitored for the depth of anaesthesia for at least 20 min with additional i.v. administration of ethyl carbamate if necessary, until they reached a stable state of anaesthesia characterized by slow, regular heart and respiration rates. Once deemed adequately anaesthetized, the rats were positioned in a stereotaxic frame with an automatic temperature control system to maintain a body temperature of 37°C. Experiments were conducted under guidelines provided by the Canadian Council for Animal Care.

Local anaesthetic lidocaine (Rafter 8, Calgary, AB) was injected subcutaneously prior to the incision of the skin above the skull. The skull was cleaned and dried, revealing reference landmark skull sutures where the reference point was taken at the intersecting point of bregma and midline sutures. Holes of ∼1 mm diameter were drilled for the placement of thalamic and cortical recording, stimulating, and injection sites (see Table 1). Bipolar stainless steel electrodes with a tip separation of ∼ 0.5 mm were used for stimulation. Single-unit and multi-unit recordings were made from glass electrodes (0.5M sodium acetate with 2% Pontamine sky blue; 2.5-6 MΩ) and tetrodes (7-8 μm platinum-tungsten wires in concentric configuration with a single centre core, ∼20 μm inter-wire distance, ∼100 μm overall diameter and impedance of 1.1-2.1 MΩ; Thomas Recording Gmbh, Germany), respectively. Intracerebral injections were made directly from an injection cannula (C315l; Plastics One, Roanoke, VA) connected to a 10 μl microsyringe lowered into the brain. Additionally, a stainless steel reference electrode was placed posterior-laterally on the right side just anterior to lambda, and a 2 × 2 mm cranial window was made above the motor or somatosensory cortices for recordings with 16 channel silicone probes (413 μm ^2^ platinum recording sites with 150 μm inter-site distance in a linear configuration; Neuronexus, Ann Arbor, MI). Paired thalamic and cortical recordings were always made on the same side, with the stimulation and injection sites also ipsilateral to recording sites.

**Table 1.** Coordinates for recording (r) and stimulating (s) electrodes. The same coordinates for the VL was also used for cannula placement.

|  | PH | VL | mCtx | VPM | sCtx(r) | sCtx (s) |
| --- | --- | --- | --- | --- | --- | --- |
| AP | 3.8 | 2.3 | 2.7 | 3.1 | 0.8 | 3.6 |
| ML | 0.2 | 1.8 | 2.0 | 2.7 | 3.0 | 2.0 |
| DV | 7.6 | variable | variable | variable | variable | 2.5 |

All recordings were pre-amplified and further amplified and filtered either with a 16-channel amplifier (A-M Systems, Carlsborg, WA) or single-channel amplifiers (Grass P511, West Warwick, RI). Regardless of amplifier differences, all signals were digitized at 250 kHz with 1-300k Hz bandpass at 1000x amplification, stored offline with DataWave hardware and software (DataWave systems, Berthoud, CO).

A typical experiment involved the lowering of the 16-channel linear probe into the cortical recording site to a depth about 2 mm below the dura. Once multi-unit activities (MUAs) from the probe had stabilized and a phase reversal of local field potential (LFP) recordings was obtained between the 4th and 7th recording site from a dorsoventral direction, the glass electrode or tetrode was lowered into the thalamic recording areas. Upon isolation of a single unit with adequate signal-to-noise ratio, or good signal-to-noise ratio on all four recording sites for tetrode recordings, at least 5 min was given before starting the experimental protocol.

### Electrical Stimulation Protocol

Once the recording had stabilized, a brief (∼1 s) PH stimulus train (0.3-0.4 mA, 100 Hz, 0.1 ms duration monophasic square waves) was delivered to elicit physiological and electrophysiological responses. This was done to ensure the associated increase in respiration with PH stimulation did not impact the stability of the recording. The stimulation protocol began after desynchronized LFP elicited by PH stimulation returned to regular UP-DOWN states (slow oscillations, <1.5 Hz; SO) seen in deep anaesthesia. The protocol began with PH stimulation at 0.3 or 0.4 mA for 10 s after 2-3 min (or up to 5 min if the firing rate of recorded units was less than 0.3 Hz) of baseline recording. Another 2 min was given to allow the LFP return to regular UP-DOWN before at least 20 0.2 ms, 0.4 mA pulses were delivered at 0.4 Hz. Following another 1-2 min recording at baseline, eight 0.1 ms pulse duration at 100Hz with 250 ms burst duration were delivered from 0.1 to 0.8 mA at 0.1 Hz (intensity steps). A wide range of parameters (0.1-0.6 mA for intensity and 0.05 to 0.5 ms pulse duration) were tested in the PH to no effect (data not shown); hence a standard 0.4 mA, 0.2 ms pulse duration stimulus (which was effective in eliciting spiking in recorded areas in the VL and sCtx) was adopted in all experiments. Finally, 0.3-0.5 mA (depending on the depth of anaesthesia and minimal current to induce cortical desynchronization) at 100 Hz with 0.1 ms pulse duration were delivered sequentially from 100 ms to 500 ms burst duration in 50 ms steps (duration step). The recording was continued for at least three more minutes before the termination of an experiment.

### Current Source Density Protocol

Current source density (CSD) was obtained through recordings from the linear 16-channel silicone probes immediately after paired thalamo-cortical recording experiments. Each recording contact was 150 μm apart and positioned through cortical layers II to VI. Stimulation electrodes were placed in the PH, ventrolateral thalamic nucleus (VL) or at layer V of the somatosensory cortex with the tips encompassing layer III to V (coordinates are provided in Table 1). One to two-minute baseline recordings were obtained before the delivery of 20 to 50 pulses at 0.3 Hz with 0.2 ms pulse width at 0.3-0.5 mA, depending on the anaesthetic level of the rats as determined in the thalamo-cortical paired recording sessions. In pilot experiments, pulse duration of 0.1 up to 0.5 ms and intensity from 0.2-0.7 mA were tested for the stimulation sites in their ability to elicit cortical spiking. However, only VL pulses appeared to be able to elicit cortical spiking; hence pulse stimulation protocols used in VL experiments were also used in somatosensory and PH experiments to allow direct comparisons between them.

### Thalamic Inactivation Protocol

The preparation of rats (n= 2) and the insertion of PH stimulating and 16-channel recording electrodes were made as described above. In addition, the injection cannula was lowered stereotaxically to the VL using coordinates in Table 1 used in CSD experiments. The experiment began with a 5 min baseline recording, followed by five 500 ms trains of 100 Hz, 0.1 ms pulses at 0.5 mA. This was repeated before the injection of 1.5 μl of saline (0.9% NaCl) or procaine hydrochloride (20% w/v in saline) was made at 1μl/min. The injection cannula remained in place for the rest of the experiment. Immediately after injection, another five bursts to the same stimulus train were delivered. From here on in, five trains of stimuli were delivered nine additional times with 5 min between each delivery.

### Data Analysis – Unit Data

All analyses of unit data were performed by using Spike2 (CED, UK) or Matlab 7.0 (Mathworks, Natwick, MA). All data files were high band passed at 500 Hz and spike sorted to separate single units from stimulation artifacts, and to isolate spike clusters from putative single units from tetrode recordings. Since the stimulation artifacts saturated the amplifier, it was possible to mark the beginning and end of each stimulation train, or single stimulation pulses by a custom-written script that detected each individual stimulus by setting an arbitrary threshold that detected only stimulus artifacts but not spikes. These markers were then used to trigger peri-stimulus time histograms in various analyses. The artifacts themselves typically lasted 2 ms.

Determination of whether a spike train increased, decreased, or did not change its firing rate was based on calculating a 95% confidence interval from 2 s bins from the 2 min baseline recordings obtained at the beginning of each experiment. If the spike rate in the 2 s immediately after the stimulus train falls outside the 95% in either direction, the unit is determined to have increased or decreased its firing rate.

For spike rate calculations for intensity and duration step protocols, again the 2 s epoch immediately after the last stimulus in a stimulus train was used as a trigger for averaging between spike trains. The same calculations were made to determine changes in spike rates during the thalamic inactivation experiment.

UP state spiking analysis was performed by using a Spike2 script (http://www.whitney.ufl.edu/BucherLab/Spike2Scripts/burstanalysis/burststat.s2s). The parameters used to isolate bursts were determined by pilot analyses, utilizing the graphical interface of the script to generate a set of parameters that best isolate what is determined as single bursts by eye. The maximum interval between spikes within a burst was set at 100 ms while the initial allowable interval between spikes at burst initiation was set at 10 ms, with a burst including a minimum of two spikes. Note these are not parameters normally associated with bursting analysis; these parameters were used to detect and generate statistics regarding the properties of MUAs during each UP state.

### Data Analysis – Field Potential Data

Analyses described here were performed by using Matlab 7.0 (Mathworks, Natwick, MA). Current source density analyses were based on 50 pulses of PH, VL, or somatosensory cortical stimulation. All pulses were delivered during SO state. Raw data were averaged over 50 pulses in a 50 ms window and 10 ms offset. The averaged waveforms were detrended first before being submitted to CSD analysis. An inverse CSD method smoothed by cubic splines was used (Pettersen et al., 2006).

Local field potential recordings were down-sampled 100x to a final sampling frequency of 250 Hz. Multi-taper spectral analyses (Mitra and Pesaran, 1999) using 1 taper with a bandwidth of 0.5 were used to calculate the power spectra from thalamic and cortical LFPs. Coherence analysis was performed with 4 tapers and a bandwidth product of 3, calculated by normalizing the squared cross-spectra from thalamic and cortical LFPs by individual spectra. The spectral and coherence ratios were derived from the differences between frequency bands between 0-10 Hz and 10-20 Hz. Since the SO to desynchronized transitions were highly stereotypical, only six pairs of LFP records from different rats were arbitrarily chosen for analysis.

### Histology

During recording sessions, constant current (50 μA, 5 min cathodal followed by 5 min anodal) was applied at the deepest recording sites. After each recording session, the rats were administered an overdose of ethyl carbamate and transcardially perfused with saline followed by 10% formalin in saline. The brains were extracted and stored in 10% formalin. Prior to sectioning, 30% sucrose solution with 10% formalin was used to cryoprotect the brain. Forty-micron slices were taken for all electrode and cannula placements, except for silicone probe recording sites, where 20 μm sections were taken instead. Electrode recording/stimulating and cannula injection sites were determined by the verification of the deepest point of each track and estimated by the depth values noted during the experiment from the microdrives.

### Statistical Analysis

All statistical analyses were performed in SPSS 15.0 (SPSS Inc., Chicago, IL) or Matlab (Mathworks, Natwick, MA). Repeated measures analysis of variance (ANOVA) was used in all statistical assessments. Polynomial contrasts were applied to determine changes of data trends over time. All reported degrees of freedom, F-statistics and *p*-values are based on Greenhouse-Geisser correction, regardless if the assumption of equal variances between groups was violated. All post-hoc comparisons are based on one-way ANOVAs, where applicable, and their statistical significance corrected by the Bonferroni method with uncorrected *p*-values presented. Some statistical tests and the generation of confidence intervals were based on surrogate data generated by bootstrapping the tested dataset. All statistical parameters generated in this fashion were at an alpha level of <.01 (or at 99%), and actual *p*-values are reported, if applicable.

## Results

### Different Thalamo-cortical Circuits Respond Differently to PH Stimulation

Data from the previous study suggested PH stimulation generally increased cortical spiking activities (Young et al., 2011b). In the motor cortex (blue line, Figure 1a), it is demonstrated that an increase of MUA spiking activity was indeed elicited upon PH stimulation from 3.84 ± 0.82 to 12.48 ± 2.60 spikes/s (F_(1, 37)_= 13.33, *p*< .001). The magnitude of the increase is more modest in the VL (1.80 ± 0.32 to 6.93 ± 0.91 spikes/s; red line in Figure 1a) due to the single unit nature of the recordings, but the detected spiking increase was statistically significant and more robust (F_(1,37)_= 34.65, *p*< .00001). The robustness of VL response to PH stimulation is corroborated by a lack of observable change in spiking variance before and after PH stimulation, whereas an increase of spike rate variability can be seen during the stimulation (Figure 1a). Contrary to the previous report, there is no evidence of spike rate increase in the somatosensory cortex (2.99 ± 0.44 to 3.81 ± 0.76 spikes/s; blue line in Figure 1b). The lack of a significant change in somatosensory spiking rate is also accompanied by a lack of significant change in VPM spiking rate (1.21 ± 0.44 to 1.95 ± 0.58 spikes/s; red line, Figure 1b).

**Figure 1.**
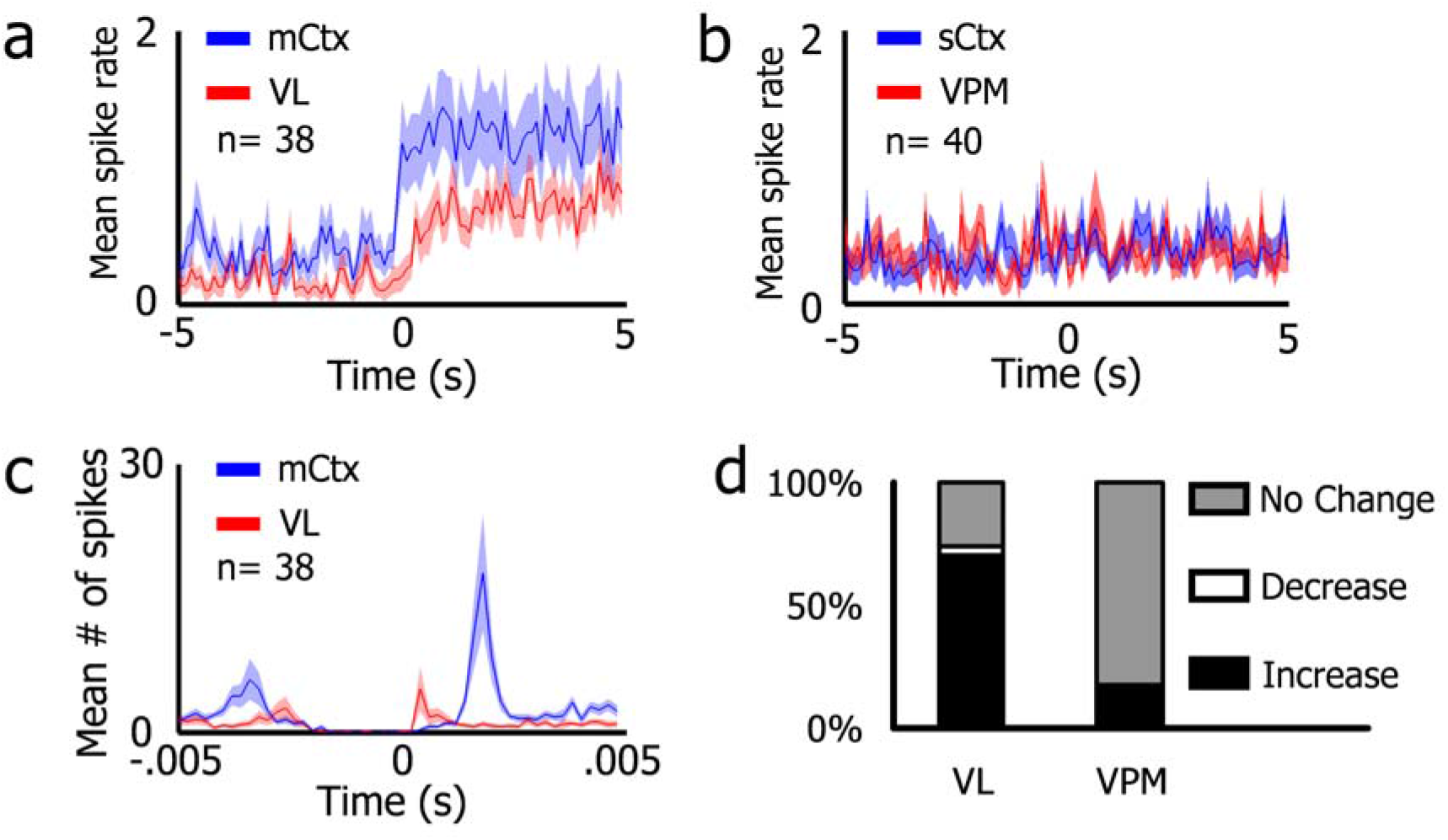
Thalamo-cortical responses to PH stimulation, (a) Paired motor thalamo-cortical (VL-mCtx) recordings reveal a rapid increase of mean firing rate from both regions elicited by continuous PH stimulation. (b) Paired sensory thalamo-cortical (VPM-sCtx) recordings did not show any marked changes in spiking rate in response to PH stimulation. (c) Peri-stimulus time histogram triggered by individual pulses within a continuous (∼10 s) 100 Hz train from the same dataset as (a). Motor thalamic spiking appears to precede motor cortical spiking. There appears to be a biphasic response where the second component shows cortical spiking leading thalamic spiking. (d) Percentage of neurons in the three response groups designated, based on the comparison of firing rates 5 s before and after the onset of PH stimulus train. More neurons increased their firing rate in the VL, whereas most did not change their firing rates in the VPM. n= number of cells.

Since an increase of motor thalamo-cortical spike rate was detected in response to PH stimulation, the stimulus-spiking relationship was further examined. A peri-stimulus time histogram based on the inter-stimulus interval of 10 ms was constructed by the triggering of each PH stimulation pulse within the train. The results indicate VL spiking occurs at 0.4 ms after the pulse, followed by the highest probability of motor cortical spiking at 1.8 ms (Figure 1c). There appear to be secondary peaks for both recording sites, with the second-highest spiking probability occurring at 7.4 ms after the pulse for the VL and 6.6 ms after the pulse for the motor cortex. Again, the differences in absolute rates described for thalamo-cortical responses are due to MUA recordings in the cortex and single units in the thalami.

### Thalamic Response May Be Dependent on PH Afferents

The response to 0.8 mA PH stimulation from all recording sites was compiled and sorted according to estimated recording depth from electrode track reconstruction. In the rostrocaudal position where VL units were recorded from (Figure 2a, right), there is a general tendency for the stimulus to elicit increases in spiking rate, from the laterodorsal thalamus extending ventrally to the most ventral aspect of the VL. This distribution corresponds to the modest PH afferent patterns to this thalamic region (Figure 2a, left).

**Figure 2.**
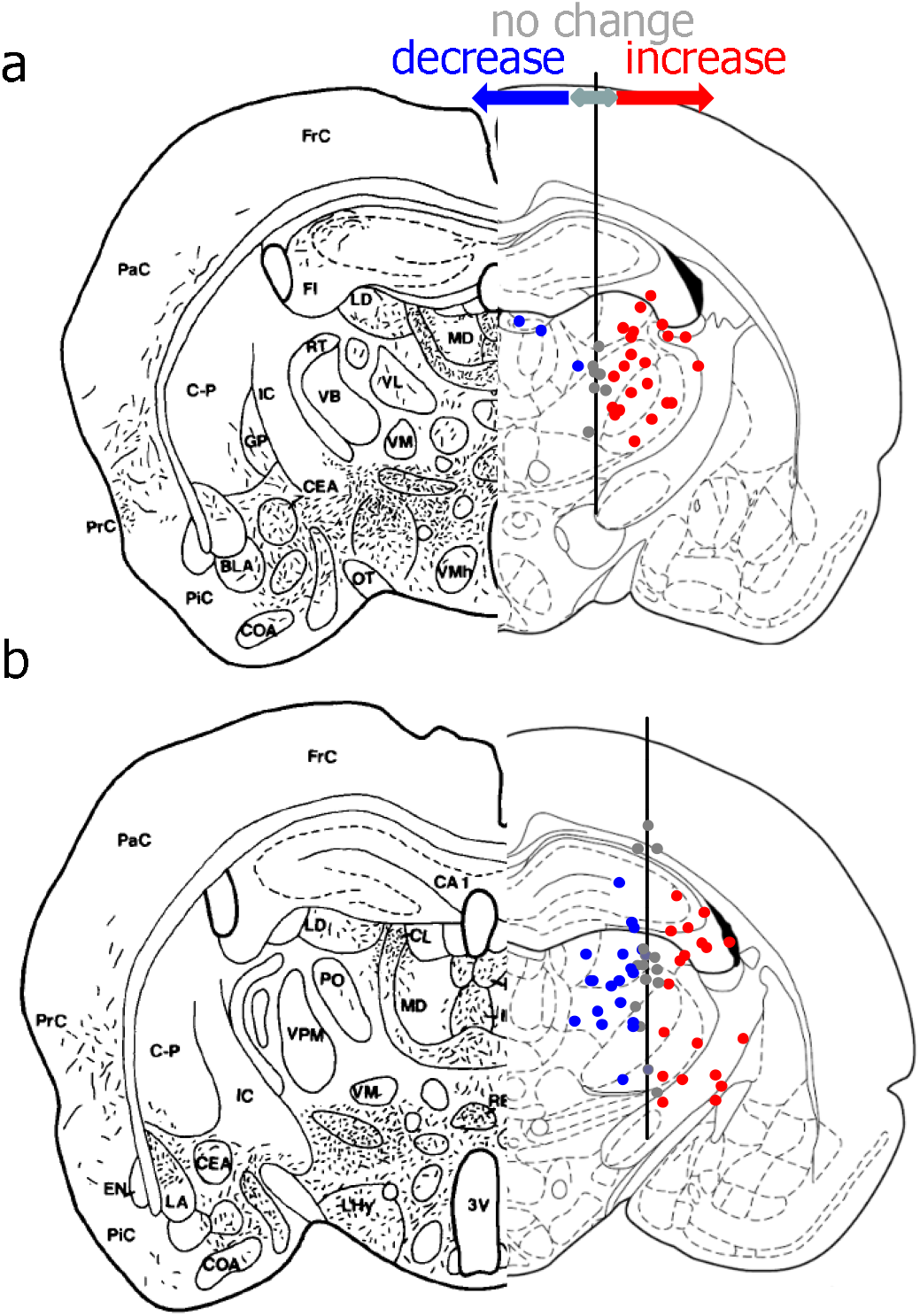
Anatomical distribution of units recorded with their responses to 0.8mA PH stimulation and the distribution of PH afferents to the area. The magnitude and direction of spike rate changes are plotted on the x-axis (increases, red; decreases, blue; and no change, grey). Recording sites are collapsed into a single medio-lateral axis (y-axis) for clearer representation of recording depth-response direction/magnitude examined here. (a) Coronal section on the right is taken from Vertes et al. (1995), illustrating an example of PH afferent fibre distribution. On the right, the positions of recorded units are plotted on the y-axis and the magnitude of their response is plotted on the x-axis, overlaying the representative rostro-caudal position of recording tracts through the VL. Most units at this level responded with excitatory responses with only a few units in the dorsal recording sites showing inhibitory responses. The majority of the units in the VL responded with excitation. (b) The same representation and organization as described in (a) but of recording sites through the VPM. There is a more even distribution of inhibitory and excitatory responses through this level, including the VPM.

As for units recorded in the region where VPM units were recorded, a more complicated pattern emerges. It can be seen from the right panel of Figure 2b that recordings from the somatosensory cortex did not significantly change their firing rates, corroborating with data already described. There was a mixed response from units recorded from the hippocampus and the laterodorsal thalamus, with a slight bias towards excitation in the latter. Then, further ventrally, the predominant response was inhibitory, with very few units demonstrating an excitatory response until passing through to the ventral half of the VPM and ventroposterior lateral thalamic (VPL) nucleus towards zona incerta and the entopeduncular nucleus. This distribution of spiking responses corresponds to the distribution of PH afferents to the area, where most of the VPM is not innervated while regions flanking dorsally (LD) and ventrally (VPL and zona incerta).

Although there is no specific demonstration of the spatial distribution of spiking response to PH stimulation to the spatial distribution of PH afferent fibres, there is a tendency for more excitatory responses from recording sites made where PH afferent fibres exist, and a more balanced distribution (perhaps biased towards inhibitory responses) where no PH afferent fibres have been reported.

### Thalamic Response to Intensity and Duration Changes of PH Stimulation

In raw traces, it can be seen that pulse protocol did not appear to elicit any observable relationship to the ongoing spiking pattern. Spiking appeared to accompany higher intensity stimulus train during intensity step protocol, whereas there appears to be increased spiking related to prolonged stimulus train during the duration stepping protocol (Figure 3a).

**Figure 3.**
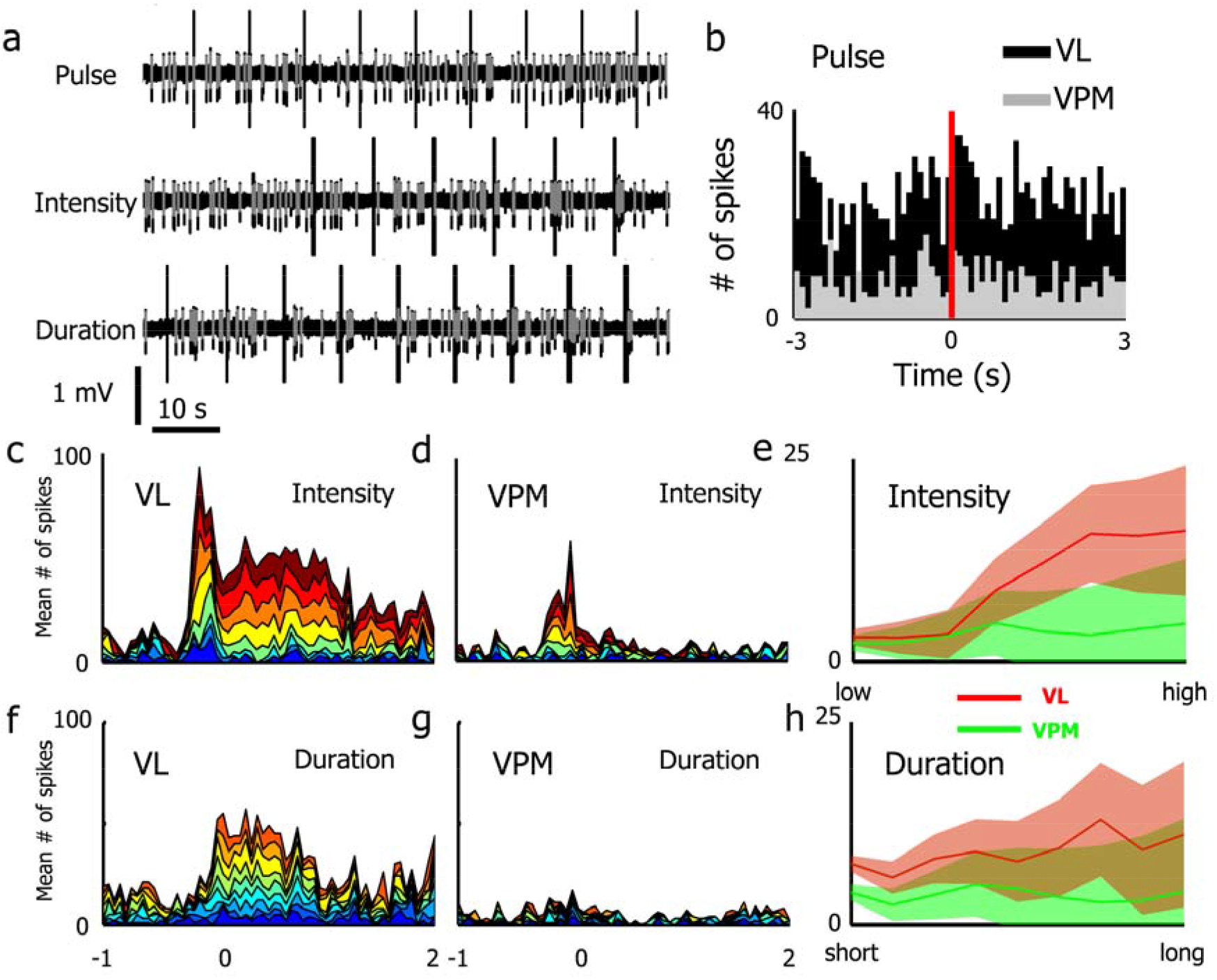
Thalamic response to various types of PH stimulation protocols. (a) Example of VL raw traces of each stimulation protocol. Grey highlights spiking activity on the black raw traces. (b) Peri-stimulus time histogram triggered by each pulse revealed no obvious rhythmicity of spiking in either thalamic nucleus using 50 ms bins. (c) Stacked representation of the number of spikes across records, where the warmest colour denotes highest PH stimulation intensity (0.8 mA) and the coolest denotes the lowest intensity (0.1 mA) used in VL recordings. Time zero marks the last stimulus delivered of the 250 ms PH stimulus train. (d) The same as in (c) but for VPM recording sites. (e) Summary of VL and VPM unit response to increasing PH stimulation intensity. The data suggest the firing rate of VL units was dependent on PH stimulation intensity, but reaches an asymptote at the highest stimulation intensity. This was not the case for VPM units. A gradual increase in the variance of firing rate can be observed in both cases. (f) VL unit responses to stimulus trains with different duration steps (100-500 ms). All trains appear to modestly increase spiking. (g) The same as in (f) but representing spiking recorded from VPM. (h) Summary of spiking elicited by PH stimulus trains of different durations. Higher duration appears to increase the amount of spiking, but this effect was weak and much variance was observed at longer trains. The gradual increase in variance is also seen in VPM recordings, without the trend to increasing spiking as a function of increased train duration.

To examine if single pulses from the PH may be able to evoke spiking responses in the thalamus, a 0.2 ms long pulse was used to stimulate the PH. There were no clear increases in spiking that followed stimulus pulse immediately (Figure 3b). However, in the VL, a number of spikes did appear to be just statistically above the overall spiking distribution over the 6 s shown (z= 1.97, *p* =.048), while no such relation was seen from VPM (z= 1.44, *p* =.15).

There is little segregation between the number of spikes recorded prior to the delivery of a 250 ms PH stimulus train (Figure 3c). Upon stimulation, there was a sharp increase of spiking, which fell slightly during the stimulus train and decreased again after the cessation of the train. Post-train spiking rate remained elevated at least up to 2 s, with the first 1.5 s showing the most obvious increase compared to baseline. The figure also illustrates that 0.4 mA appeared to elicit significant increases in spiking, while the areas under the curve for the last two intensities did not seem to differ.

Figure 3e summarizes the above observations by plotting the mean number of spikes generated in each 2 s epoch immediately after the stimulus train. Indeed, a linear increase of spiking can be observed in the VL but not VPM (F_(1,18)_= 19.12, *p*< .0005 and F_(1,18)_= 1.11, *p*= .31, for respective linear trends). However, the increase in spiking observed in VL reached an asymptote for the two highest stimulation intensities, suggesting the majority of units can be maximally driven at 0.7 mA (cubic trend, F_(1,18)_= 7.29, *p*< .05). In addition, there appears to be a progressive increase of variance as a function of increasing stimulation intensity in both the VL and VPM, suggesting the intensity-spiking relationship in some units may be relatively non-linear and/or the magnitude of response to each intensity step is different among units recorded (i.e. the intensity-spiking function slopes are different).

All duration step stimulus trains were delivered at 0.3-0.5 mA, depending on the minimal current step that elicited cortical desynchronization. Similar to responses elicited by intensity steps, the first two duration steps (100 and 150 ms train duration) did not appear to evoke any spiking in the VL during the train or post-stimulus (Figure 3f). There was no evidence for longer trains to evoke longer periods of persistent firing upon stimulus train termination. There did not appear to be any differences between firing rates before or after the stimulus trains at any duration step in the VPM, although some evidence prolonging spiking state can be observed during and immediately after a stimulus train (Figure 3g). Figure 3h summarises the VL and VPM responses to PH duration step stimulus trains, where VL showed linear increases of spiking activity as a function of duration step (F_(1,18)_= 10.13, *p*< .01) and VPM showed no changes (F_(1.98,37.52)_= 2.63, *p*= .09), but an increase in response variability similar to what was described for intensity step experiments can be seen from both areas.

### PH Stimulation Elicits Similar Changes in Thalamo-Cortical LFPs

Raw recordings from the motor and somatosensory thalamo-cortical pairs are illustrated in Figure 4a. Clear, larger amplitude, regular activities can be seen prior to PH stimulation and small amplitude irregular activities can be seen following the stimulus train characterized by saturated stimulus artifacts. These changes are reflected in the power spectra of LFPs, where a reduction of slow frequency activities was coupled by an increase in >10 Hz activities in both the VL and VPM (Figures 4c and d, respectively). These changes are more prominent in the thalamus as thalamic SO was of smaller amplitude. When this state transition is calculated as a proportion of slow and fast activities found in the LFP, it is clear that PH stimulation presumably evokes a more activated state based on the transition from regular UP-DOWN states to persistent small-amplitude irregular activity seen in awake animals in both the cortex and the thalamus (Figure 4d). Spectral coherence indicates a reduction in the slow frequencies and an increase in higher frequencies in both motor and somatosensory thalamo-cortical pairs (Figures 4e and f, respectively). This change appears to be stronger in the somatosensory thalamo-cortical pairs. Finally, corroborating power spectral and coherence findings discussed above, the state change PH stimulation invokes similar changes in the spectral content of LFPs and demonstrates coupling of SO to desynchronized LFP transition (Figure 4g).

**Figure 4.**
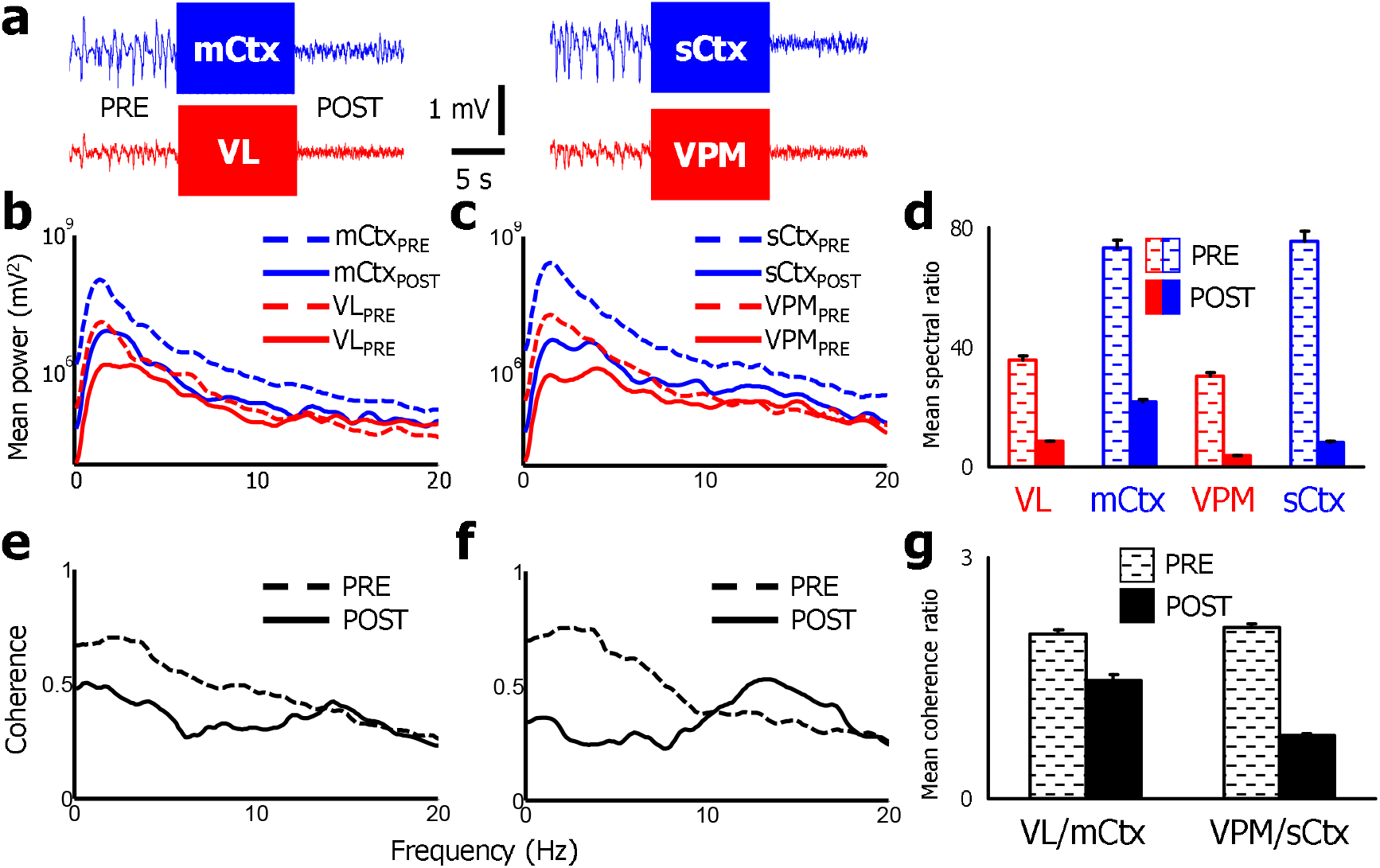
Spectral changes before and after PH stimulation train. (a) Raw LFPs from the motor (left) and sensory (right) thalamo-cortical recordings. Clear transitions from large amplitude, semi-regular SO to smaller irregular activities can be seen after the stimulus train, which produced characteristic amplifier saturation. (b) Power spectral changes of motor thalamo-cortical LFPs indicate a decrease of power at ∼2 Hz range after PH stimulation, coupled with a slight increase of activities at >12 Hz. (c) The same as in (b) but representing spectral changes from the somatosensory thalamus and cortex. Similar changes as described for their motor counterparts are shown. (d) Quantitative summary of spectral changes elicited by PH stimulation, corroborating the effect of PH stimulation in suppressing SO and increased higher frequency activities in the 5 s post-train period. (e) Coherence spectra for the motor thalamo-cortical LFPs. The change in coherence mirrors those reported for power spectra, where a decrease in lower frequencies was matched by an increase of coherence in higher (> 7 Hz) frequencies. (f) The changes described for motor thalamo-cortical recordings are also seen in their somatosensory counterparts, with a clearer dissociation of low vs. high frequency components before and after stimulation. (g) Corroborating with the trends described above, there was a significant shift of coherence from lower frequencies to higher frequencies, with the effect being more prominent in paired somatosensory thalamo-cortical recordings.

### CSD Profiles of Thalamic, Cortical, and PH Pulse Stimulation

Pulse stimulations of the VL evoked an early source in the border of layers V and VI, which appears to propagate further ventrally and clearly propagates dorsally to layer V, flanking the prominent sink that arises in layer VI (Figure 5a, top). More superficially, a small sink can be seen in dorsal layer V before two additional sources in layers II/III and I follow the deeper layer source with a short latency (∼3 ms). Raw traces below the CSD profile (Figure 5a, bottom) indicates the source at the layer III and V boundary may be more concurrent than being separated by a small latency (green and red lines) and that VL stimulation evoked spiking is highly reproducible, appearing in over 50 averaged waveforms.

**Figure 5.**
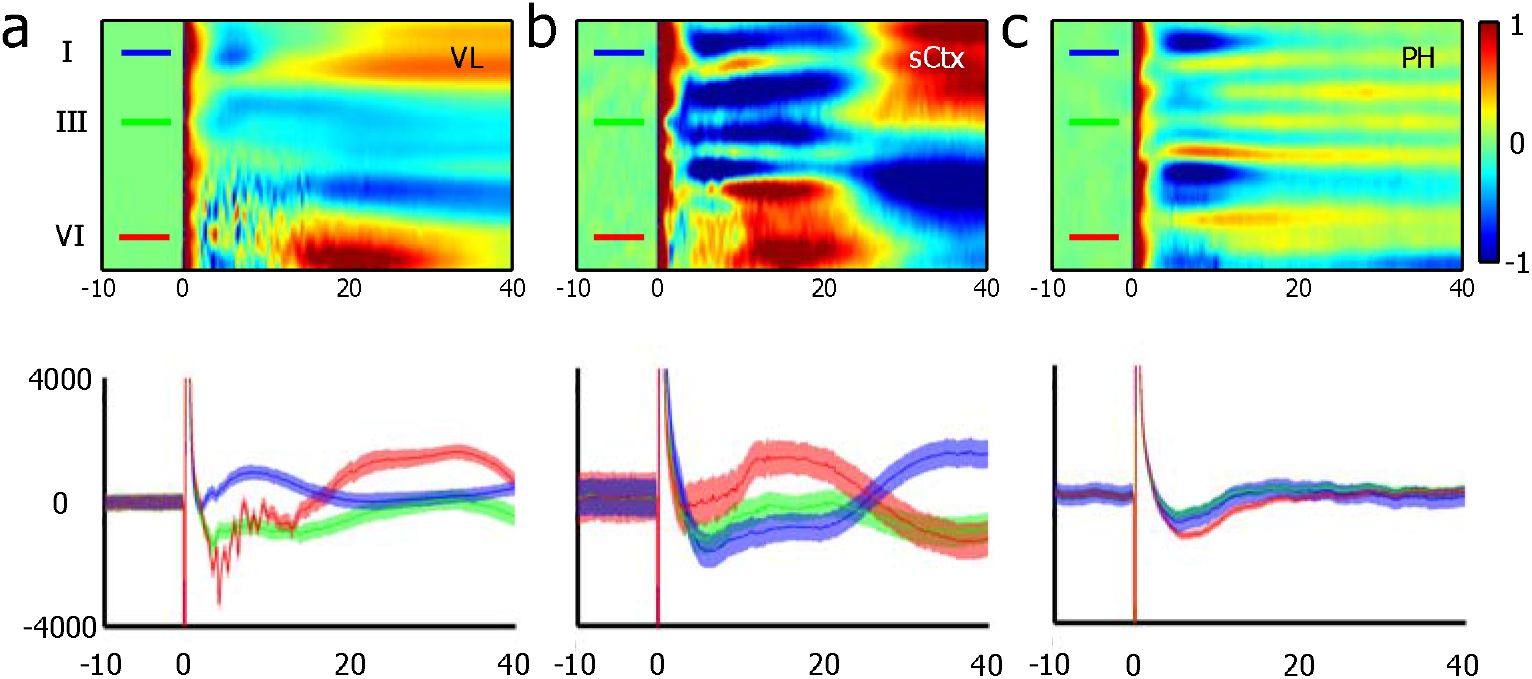
Current source density profiles from different stimulation sites. (a) Current source density plot (top) from VL stimulation pulses. The largest and earliest source/sink (cool/warm colours) pair is observed in more ventral recording sites and smaller source/sink pairs can also be observed in more superficial layers. Normalized raw traces from different layers are plotted below the CSD profiles, showing the relative sizes of evoked potentials. (b) Current source density profile of the motor cortex elicited from somatosensory cortical (sCtx) pulse stimulation. Differing from (a), most prominent and earliest sources are located in the more superficial layers. A deep (layer VI) source-sink pair that seem to emerge as early as the more prominent and superficial one found in layer III, is corroborated by the select raw traces displayed below. (c) Current source density profile elicited by PH stimulation yielded a similar pattern to those observed in somatosensory cortex stimulation in the first 10 ms. It is unclear if the profile was indeed evoked, given the lack of any late component after initial differences in response phase polarity (see also raw traces below).

Somatosensory cortex stimulation resulted in a large source predominantly in the intermediate (layers II/III) in the motor cortex, with the earliest response seen in layer II/III, and other prominent sources following 2-3 ms later in dorsal layer V and in layer I. Interestingly, it appears a brief layer VI source also coincides with the primary source in layer II/III. Regardless, the primary response in layer V/VI is associated with sink currents that last ∼10 ms before giving away to a source coupling the large sink in more superficial layers (Figure 5b, top). The raw waveform averages presented below corroborates the CSD findings, and indicate that evoked responses were perhaps more variable in comparison to other stimulation protocols (Figure 5b, bottom; c.f. Figures 5a and c bottom panels).

With PH stimulation pulses, there did not appear to be differences in latency between any current sources or sinks in the motor cortex. Multiple source/sink pairs exist, with the layer V source being the most prominent, followed by another source in layer I, then in layer VI (Figure 5c, top). There are many smaller source/sink pairs throughout the motor cortex, but are most likely artifacts due to the lack of an actual evoked response indicated by the lack of change over time, resulting in the representation of differences in residual direct components of the LFPs. Raw averaged waveforms support this interpretation as sampled responses from layers I, III, and VI all appear similar, with no late components visible (Figure 5c, bottom).

### 5.3.6 Thalamic Inactivation Abolishes Cortical Spiking Elicited by PH Stimulation

Prior to procaine infusion (Figure 6a, left) and in saline trials (data not shown), PH stimulus trains readily evoked spiking during and immediately after each train with associated changes from SO to desynchronized LFP states. The rate histograms (Figure 6a, grey trace) suggest high-density spiking behaviour during each UP state were transformed into persistent spiking during desynchronized LFP state. Immediately after procaine infusion (Figure 6b), PH stimulus trains still elicited cortical desynchronized states with little spiking during the train itself and an almost complete lack of spiking post-stimulus. This effect of thalamic inactivation appears to be selective, as clustered spiking during cortical UP states remains present.

**Figure 6.**
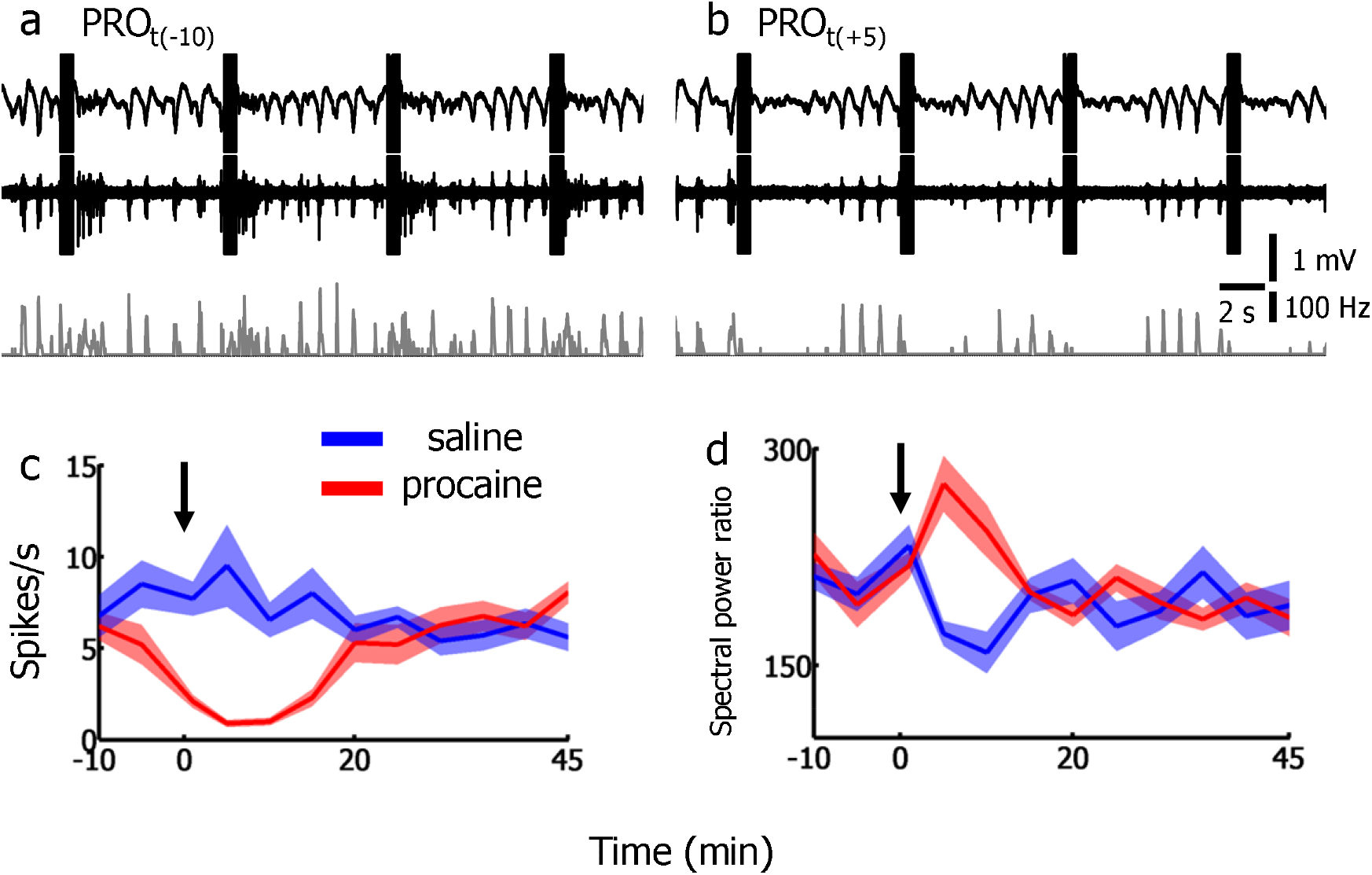
Procaine injection to the VL inhibits spiking in the motor cortex elicited by PH stimulation. (a) Raw traces, accompanied by a firing rate plot obtained from the motor cortex prior to procaine injection. Spiking activity can be seen clearly following PH stimulation trains. Rate histograms also suggest a switch from clustered spiking from MUA activity during cortical LFP UP states to persistent firing during brief desynchronization states. (b) Cortical desynchronization occurs after PH stimulation trains in the absence of spiking activity after procaine injection. (c) Time-course of spike rate changes during experiments. The downward pointing arrow at time zero indicates the first five stimulation trains delivered right after procaine infusion. Thalamic procaine injection induced a clear decrease of post-train spiking rates, with recovery at the 20 min mark. (d) Spectral power ratio between slow (0-10 Hz) and fast (12-20 Hz) activities during the 2 s after each stimulus train were increased by procaine injection to the thalamus. The changes mirror the time course seen in spiking data, falling to baseline 20 min after injection.

Comparisons between spiking during and after each of the five stimulus trains delivered at each time point revealed no change in spiking rate (F_(3.45,65.55)_= 1.85, *p*= .14) throughout the experiment (Figure 6b, blue line). In contrast, there was a statistically significant change in spiking rate with procaine infusion experiments (F_(3.06,58.14)_= 17.31, *p*< .0001). Specifically, a prominent cubic function (F_(1,19)_= 18.03, *p*< .001) approximates the change of spiking rates with decreases coinciding with the time of procaine injection (Figure 6b, red line). As a function of time, there were statistically significant changes in the slow/fast LFP activity ratio with both saline and procaine injections (F_(2,6.01)_= 20.51, p< .01 and F_(1.3,3.89)_= 17.43, p< .05, respectively). Polynomial contrasts suggest both conditions show prominent higher-order polynomial functions in addition to an expected quadratic function representing drug effect and recovery (see Table 2 for a summary).

**Table 2.** List of polynomial contrasts for slow vs. fast LFP ratios recorded in the motor cortex based on 2 s post-stimulus epochs. Corrected statistically significant trends are bolded.

|  | SAL |  | PRO |  |
| --- | --- | --- | --- | --- |
|  | F-ratio | *p*-value | F-ratio | *p*-value |
| linear | 2.09 | 0.24 | 9.94 | 0.05 |
| quadratic | 13.36 | 0.04 | 25.41 | 0.02 |
| cubic | 229.74 | **0.00** | 65.03 | **0.00** |
| 4th order | 25.35 | 0.02 | 0.11 | 0.76 |
| 5th order | 9.67 | 0.05 | 50.36 | 0.01 |
| 6th order | 6.04 | 0.09 | 67.91 | **0.00** |
| 7th order | 26.86 | 0.01 | 16.20 | 0.03 |
| 9th order | 315.61 | **0.00** | 11.47 | 0.04 |
| 10th order | 7.05 | 0.08 | 17.82 | 0.02 |
| 11th order | 1.12 | 0.37 | 0.53 | 0.52 |
| 12th order | 2.13 | 0.24 | 5.37 | 0.10 |

## Discussion

This study examined the responses of two thalamo-cortical circuitries to different types of PH stimulation. Preferential activation of the motor thalamo-cortical circuitry over somatosensory thalamo-cortical circuitry was demonstrated, possibly due to the relative amount of PH innervations to the respective areas. By varying stimulus strength and duration, it was determined that the PH does not directly elicit spiking in the cortex, and there is an intensity-dependent function of post-stimulus PH spiking rate. These epochs of increased spiking in the motor circuitry are associated with coherent changes in LFPs, where large amplitude SO is transformed into fast, small amplitude desynchronized activities. The lack of a direct PH involvement in increased motor cortex excitability is supported by dissimilarities in CSD profile differences. Finally, it was revealed that VL inactivation can abolish PH-elicited spiking in the motor cortex, suggesting that observed cortical excitation is mediated by thalamic relay. There is also evidence for disruption of thalamo-cortical dynamics during the characteristic UP-DOWN states under anaesthesia, supporting thalamic involvement in the expression of spontaneous UP-DOWN states in the cortex.

Multiple lines of evidence reported here suggest that at least a part of PH’s facilitatory effects on motor behaviours may be mediated by its ability to evoke spiking activity in the motor thalamus and cortex. Particularly, the non-stereotypical nature of motor behaviours elicited by PH stimulation may be due to the activation of higher-order areas such as the motor thalamo-cortical circuitry. Similar findings have been reported in the previous study (Young et al., 2011b), where PH stimulation was found to induce increased cortical spiking associated with desynchronized LFPs. However, this study found no evidence for PH stimulation to increase the excitability of the somatosensory cortex, which was reported in the previous study. Differences in recording sites, and the fact that data from other cortical areas such as the parietal and cingulate cortices were reported together with the somatosensory cortex may have masked its lack of spiking response from the somatosensory cortex to PH stimulation.

The idea that PH may selectively enhance spiking in the motor cortex through the VL is supported by the coincidental increases in both motor cortex and VL spiking rates during and after PH stimulation. These increases are coupled with concurrent LFP state changes and maximal increases in coherence at the 12-15 Hz band. Of course, given the reciprocal and highly interactive nature of thalamic and cortical circuitries, it is not possible to infer the direction of driving based on these data. However, the demonstration of: 1) thalamic spiking occurring before cortical spiking in response to PH stimulation; 2) a weak but significant bias of firing probability immediately after a PH pulse in VL but not the VPM and; 3) similarities in the location of PH and VL evoked sources and sinks all suggest the increase of spiking in the motor cortex is relayed through the VL. Crucially, VL inactivation through procaine infusion virtually abolished cortical spiking triggered by PH stimulation during and after PH stimulation, and its recovery to baseline levels as well as the lack of an effect of saline infusion are supportive of the conclusion that PH mediates motor cortical excitation through the VL. Paired ensemble recordings concurrent with local inactivation are necessary to unequivocally verify whether the lack of PH-elicited cortical spiking is due to the silencing of thalamic neurons. Finally, the lack of evoked spiking and strong current sources and sinks in the motor cortex by PH stimulation pulses argue against activity propagation in a cortico-thalamic direction despite the existence of anatomical substrates to support such propagation.

The argument that PH selectively drives the VL and subsequently drives the motor cortex is also supported by the anatomical distribution of recorded units through the VL and VPM that demonstrated an excitatory response. Specifically, the number of excitatory responses was higher in the VL compared to the VPM, where only the most dorsal and ventral aspects of the nucleus had higher probabilities of being excited by PH stimulation. This roughly corresponded to the PH afferent fibre distributions in the respective regions, where the VL receives modest innervations but the VPM is virtually devoid of PH afferents. Extending from the VL and VPM, overlaying laterodorsal thalamic nucleus also receives generous innervations from the PH, more so in its medial than its lateral aspect. This is also correlated with LD recording sites through to the VL and VPM, where more medial recordings through the VL yielded a high number of neurons excited by PH stimulation and a more even excitatory/inhibitory distribution at the level of VPM. In fact, the reported increases in cortical spiking in the previous study may be mediated by the excitatory effects of PH stimulation on the cingulate and/or parietal cortices through the LD, through similar mechanisms as described here for VL and motor cortex. Finally, PH also projects densely to areas ventral to the VPM such as parts of the VPL and zone incerta, both exhibiting a predominant excitatory response upon PH stimulation.

The involvement of the thalamus on cortical desynchronization has been mostly attributed to the intralaminar nuclei (Steriade, 1970), with larger contributions from the basal forebrain (Buzsaki et al., 1988; Jones, 2003, 2004) and neuromodulator centres in the brainstem (Dringenberg and Vanderwolf, 1997, 1998). All these areas receive extensive innervations from the PH (Vertes et al., 1995; Vertes and Crane, 1996). It is demonstrated here that VL inactivation does not abolish cortical desynchronization but nevertheless may have an effect on the duration or frequency content of PH-elicited cortical desynchronization. It is not possible with the current information to determine if both duration and frequency content are involved in this detected change due to the use of time-fixed epoch data for the analyses. Since saline injection also produced a modest effect (i.e. induced a decrease in the ratio, suggesting more faster and/or less slower activities in the given epochs) on cortical desynchronization, it is unclear if statistically significant changes detected here are representative of underlying cellular processes and what its biological mechanisms could be.

Cortical UP-DOWN states are known to be reflected in the thalamus (Dossi et al., 1992; Steriade et al., 1993b; Amzica and Steriade, 1995; Steriade et al., 1996; Timofeev and Steriade, 1996), but a series of *in vivo* and *in vitro* studies have suggested cortical UP-DOWN states are generated within the cortex itself (Steriade et al., 1993a; Sanchez-Vives and McCormick, 2000; Compte et al., 2003), possibly by the intrinsically bursting neurons in layer V (Silva et al., 1991; Wang and McCormick, 1993; Amzica and Steriade, 1998; Sanchez-Vives and McCormick, 2000), propagated in a rostrocaudal direction independent of the thalamus (Sanchez-Vives and McCormick, 2000; Massimini et al., 2004; Massimini et al., 2007). Despite the perceived lack of importance of thalamic involvement in the expression of the cortical UP-DOWN states, many studies have demonstrated that indeed the thalamus at least modulates the expression of normal cortical UP-DOWN states (Cossart et al., 2003; Shu et al., 2003; MacLean et al., 2005; Rigas and Castro-Alamancos, 2007) and some thalamic neurons may spontaneously exhibit UP-DOWN states without the cortex (Williams et al., 1997; Hughes et al., 1999; Hughes et al., 2004; Blethyn et al., 2006; Zhu et al., 2006). This study adds to the accumulating evidence for thalamic involvement in cortical UP-DOWN states by demonstrating: 1) an increase in spike numbers per each UP state; 2) briefer windows of spiking during an UP state and; 3) increased inter-UP state interval and the length of UP-DOWN state cycles of motor cortical activities in VL procaine infusion experiments. However, only the changes in inter-UP state interval appear to have a procaine-dependent effect, based on the similarity of its time-course to PH-elicited cortical spike rates, and the assumption that the ability to elicit cortical spikes by PH stimulation reflects the absolute influence of procaine on the VL. Additional changes during saline infusion experiments suggest much variance exists in spontaneous UP-DOWN state properties. Again, concurrent paired ensemble recordings from the motor cortex and VL with procaine infusion may provide insights to if and how VL inactivation may lead to changes in spiking characteristics during spontaneous UP-DOWN states.

Data from the present study suggest electrical stimulation of the PH selectively excites VL, most likely through its presumed direct glutamatergic projections to the area, and then, in turn, excites the motor cortex. This excitatory PH-thalamo-cortical axis described here may be the cellular substrate for the ability of PH stimulation to elicit non-stereotypical motor behaviours. Selective augmentation of this circuit may account for, at least in part, the therapeutic effects of PH stimulation in parkinsonian akinesia (Jackson et al., 2008; Young et al., 2009; Young et al., 2011a). Paired thalamic and cortical recordings and inactivation experiments in the freely moving animal would provide much-needed relevance for the excitatory PH-thalamo-cortical pathway described here.

## Acknowledgements

This research was supported by Natural Sciences and Engineering Research Council (NSERC) research grant to BHB (A9935). CKY was supported by Parkinson Society Canada (PSC) and Parkinson Society of Southern Alberta (PSSA).

## Contributions

CKY conceived the study, carried out all experiments and analyses. BHB provided supervision and secured funding. CKY prepared the manuscript and both CKY and BHB reviewed and edited the manuscript

